# Stochasticity-induced stabilization in ecology and evolution

**DOI:** 10.1101/725341

**Authors:** Antony Dean, Nadav M. Shnerb

## Abstract

The ability of random environmental variation to stabilize competitor coexistence was pointed out long ago and, in recent years, has received considerable attention. Here we suggest a novel and generic synthesis of stochasticity-induced stabilization (SIS) phenomena. The storage effect in the lottery model, together with other well-known examples drawn from population genetics, microbiology and ecology, are placed together, reviewed, and explained within a clear, coherent and transparent theoretical framework. Implementing the diffusion approximation we show that in all these systems (including discrete and continuous dynamics, with overlapping and non-overlapping generations) the ratio between the expected growth and its variance governs both qualitative and quantitative features of persistence and invasibility. We further clarify the relationships between bet-hedging strategies, generation time and SIS, study the dynamics of extinction when SIS fails and the explain effects of species richness and asymmetric competition on the stabilizing mechanism.

## I. INTRODUCTION

Studies in ecological genetics have documented temporal fluctuations in fitness among alleles in many species [4, 6, 14, 22, 24, 41, 42, 46] and long-term ecological field studies have documented fluctuations in the recruitment and death components of Malthusian parameters in many others [4, 5, 13, 30, 32, 33, 39]. However the extent to which this variability affects the long-term outcome of competition and biodiversity is not well understood. One obvious reason is that even long-term field studies offer but a fleeting glimpse into ecological and evolutionary processes that play out across millennia.

Theory partially fills this gap in knowledge by providing means to explore long-term outcomes. Typically, a dynamical model is proposed with variable parameters that have been calibrated empirically over a relatively short time scale [23, 48, 49]. The model is then analyzed, numerically or analytically, to determine the conditions under which fluctuations in competition promote (or destabilize) biodiversity, to assess the impact that such fluctuations have on rates of allelic/species turnover and to evaluate the likelihood that new alleles/species will invade and become established in the population/community (from hereon “species” in a “community” is synonymous with “alleles” in a “haploid population”). Increasing attention is also being paid to models of fluctuating competition to determine if their long-term predictions are consistent with field observations [5].

Most of the literature in the field, particularly a series of papers that laid the foundations of the modern coexistence theory [8–10], focuses on rare species dynamics along the log-abundance (or log-frequency) axis. Emphasis is placed on the **logarithmic** mean growth rate when rare, 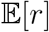 as a measure of invasibility. Here we suggest an alternative approach, based on the ratio between the expected growth when rare *μ* and its variance *g*, when both parameters are measured along the **arithmetic** abundance (or frequency) axis. In what follows we will claim that our *μ/g* analysis is better than 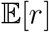-based methods for two reasons:

1. To calculate 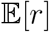 one considers the dynamics of a rare species along the log-abundance axis. This choice is natural when the growth rate is a linear function of abundance [*dx*/*dt* = *r*(*t*)*x*] as the dynamics on the log-abundance scale is a simple random walk. This choice is unnatural when growth rates are not linearly dependent on abundance; for an Allee effect or a recessive allele the growth rates are quadratic functions [*dx*/*dt* =*r*(*t*)*x*^2^]. The *μ/g* analysis presented here is more transparent, more flexible, and allows for a synthesis of different cases within a common simple mathematical framework.
2. Our *μ/g* analysis is a better indicator of invasibility and persistence. As explained in Appendix **I**, 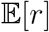 is a qualitative, but not a quantitative, indicator. While a change in the **sign** of 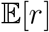 reflects a substantial change in invasibility, the **magnitude** of 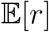 is less important; “a bigger 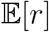” implies neither greater invasibility nor longer persistence. Our parameter *μ/g* changes sign when 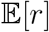 changes sign, while its magnitude dictates the likelihood of invasion and the time to extinction. Unlike 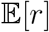, both the sign and the magnitude of *μ/g* are directly related to persistence, such that an increase in the numerical value of *μ/g* indicates greater stochasticity-induced stabilization (SIS).

To facilitate discussion, we restrict our analyses to a parameter regime in which abundance variations are not large and the diffusion approximation [15, 35] holds. Our theory is focused on a large, well-mixed community. Effects of demographic stochasticity or the possible impact of spatial structure are assumed to be small and are discussed only rarely.

Armed with our *μ/g* approach, we explore, in sections II and III, the means by which fluctuating selection promotes coexistence in four models: the lottery model of Chesson & Warner [12], the serial transfer model of Yi & Dean [51], a diploid genetic model with complete dominance by Haldane & Jayakar [26], and the Moran/chemostat models of Shnerb [16–18, 29, 43] and Dean [19–21] and their collaborators. Our focus is on competition between two alleles/species in which the amplitude of selection depends on the environmental conditions. For simplicity, and without loss of generality, we allow the selection coefficients to assume either of two values (dichotomous, or telegraphic, stochasticity). *Mutatis mutandis*, our analysis is relevant to game theory where different strategies are considered as alleles or species.

In section IV we clarify the relationships between SIS and bet-hedging mechanisms like adult longevity, seed bank and so on. Contrary to the prevailing view according to which these mechanisms are essential to SIS, we show that their main effect - to reduce the effective amplitude of environmental variations - impedes SIS. If bet-hedging mechanisms improve stochasticity induced stability, it is probably just because they tend to lengthen the generation time.

Section V is devoted to the usage of *μ/g* as a quantitative measure of stability. We show that both the probability of invasion (Eq. S8) and the persistence time (Eq. 49) are directly related to *μ/g*. In that context, the relationships between our *μ/g* and the parameter used in modern coexistence theory, 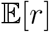, are clarified in Appendix I, where we explain in some detail why *μ/g* a more suitable diagnostic tool. When the SIS mechanism fails the system reaches extinction through a very interesting dynamics that involves rejuvenation bursts, this process also is described in section V. Finally, in section VI we explain how the SIS effect weakens in diverse communities and in section VII we consider the ability of stochasticity to induce stability in asymmetric scenarios where one of the species is fitter than the other.

## II. *μ/g* ANALYSIS IN THE ARITHMETIC FREQUENCY DOMAIN

Consider an arbitrary zero-sum two-species/allele problem in an infinite community/population for which the frequency of the focal species is *x_t_* at time *t*. Let the dynamics of *x* be described by a map that gives *x*_*t*+*δ*_ in terms of *x_t_* and other parameters, including the state of the environment *E*. Analyzing such maps is standard practice in population genetics, where *δ* is the generation time and the generations are non-overlapping. Here, we shall think of *δ* as the *persistence time of the environment* (i.e., the typical time for which the environmental conditions remain constant). In what follows we provide examples in which this persistence time is shorter, longer and also equal to the generation time.

To understand stochasticity induced stabilization, consider a single step of this map from *x*_0_ to *x_δ_*. We now omit the index *δ* and speak about *x*_1_ as a function of *x*_0_ and *E*, the prevailing environmental conditions. Accordingly,

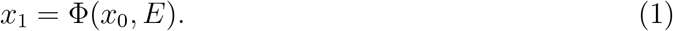

If the environment is fixed (i.e. if there is no environmental stochasticity) then *E* is fixed and for every *m* the map *x*_*m*+1_ = Φ(*x_m_, E*) has the same parameter *E*. A simple map familiar from population genetics gives the focal allele frequency in the next generation as

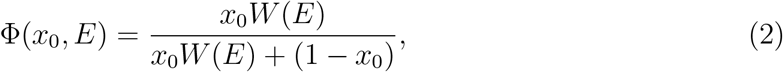

where *W*(*E*) is relative fitness.

In a closed system (with no migration and no speciation/mutation) Φ must satisfy two conditions:

1. Φ(0, *E*) = 0, meaning that the dynamics halts at *x*_0_ = 0, the extinction point.
2. Φ(1, *E*) = 1, so the dynamics halts at *x* = 1, the fixation point. Coexistence is *stochasticity-induced* if it does not occur under fixed environmental conditions. Accordingly, we require,
3. For any *fixed E*, either Φ(*x,E*) > *x* for 0 < *x* < 1 or Φ(*x, E*) < *x* for 0 < *x* < 1. This condition is *sufficient* to ensure that under the map (1) the value of x either shrinks to zero or grows to one.

In Figure 1 (which refers to the another map, given in Eq. (16) below) the red and the blue curves represent maps that satisfy conditions (1-3). Undertheredmap *x*_*m*+1_ < *x_m_* and the frequency shrinks to zero. Under the blue map *x*_*m*+1_ > *x_m_* and the frequency grows to one.

**FIG. 1:**
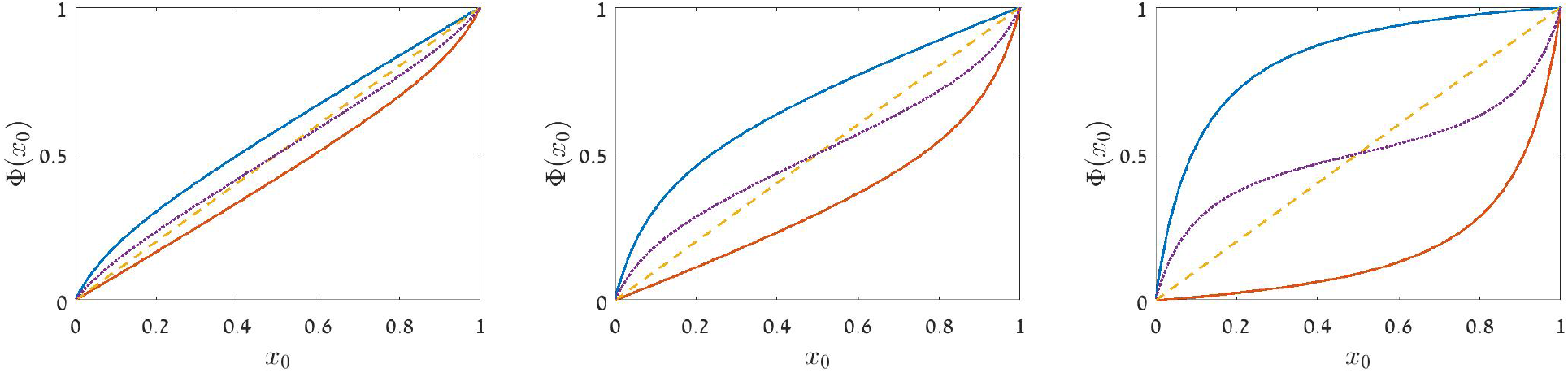
The lottery map, Φ(*x*_0_, *E*) in Eq. (16), is plotted against *x*_0_ for *δ* = 0.2 (left), *δ* = 0.5 (middle) and *δ* = 1 (right). Crossing of the dashed yellow line *x*_1_ = *x*_0_ corresponds to the fixed points of the map where Φ(*x*_01_, *E*) = *x*_0_. Above the yellow line the frequency of the focal species grows, below this line the frequency of the focal species shrinks. Φ(*x*_0_, 10) (blue line) always lies above the dashed yellow diagonal and so the focal species sweeps to fixation. Φ(*x*_0_, 0.1) (red line) always lies below the dashed yellow diagonal and so the focal species goes extinct. The mean map, (Φ(*x*_0_, 10) + Φ(*x*_0_, 0.1))/2 is represented by the dotted purple line. The expected growth of each species is positive when each is rare and the mean map has an attractive fixed point at *x*_0_ = 0.5. These figures seem to suggest that SIS becomes stronger (higher asymmetry between curves) as δ increases. **This is wrong**: diffusive trapping causes SIS to decrease with *δ* and vanish when *δ* = 1, as demonstrated in Fig. 2.

What happens when the environment changes its state, for example, when *x*_*m*+1_ = Φ(*x_m_, E_m_*) and *E_m_* is a given stochastic process? It turns out that changes in *E_m_* promote coexistence, defined here as the ability of each species to invade the system when rare, if two criteria are fulfilled.

First, each species must have a **positive expected growth when rare** (*x* ≪ 1). If the expected growth (the average taken over all the states of the environment) is negative, then the expected frequency of the species decays with time and so it cannot invade.

Second, each species must avoid **diffusive trapping when rare**. A positive expected growth is **not** sufficient for invasion. When the expected growth is zero, the dynamics is controlled by random abundance variations and the system performs some sort of “random walk” along the *x* axis. In this random walk the step size approaches zero as *x* → 0. Once the abundance reaches the extinction zone (close to zero) its dynamics slows down tremendously. As a result, when the expected growth is zero the probability density accumulates closer and closer to the extinction point, a process that we analyze in detail in section V below. Persistence is ensured only if the expected growth is strong enough to overcome this effect.

Accordingly, a system that satisfies conditions 1-3 produces SIS whenever the expected growth when rare is positive for both species and there is no diffusive trapping. Let us explain the conditions under which a map like (1) satisfies these two requirements.

### A. Positive expected growth when rare

The frequency of the focal species when rare (*x* ≪ 1) is,

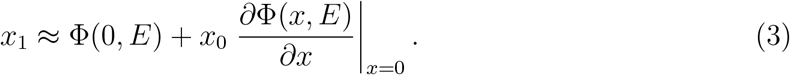

Condition (1) above ensures that the first term vanishes and so the focal species will grow on average if,

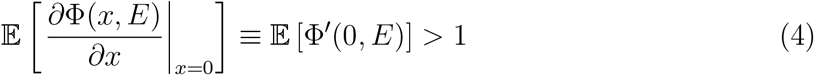

where 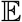 is the expectation (average) taken over all possible environments *E* (given their probabilities) for a fixed value of *x*. On the other hand, when *x*_0_ is close to one,

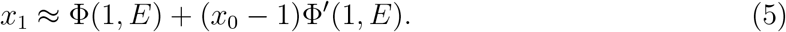

Condition (2) implies that Φ(1,*E*) = 1 so

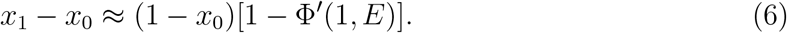

and the focal species cannot fix (i.e., its rival species grows when rare) if,

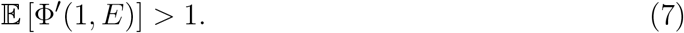

To conclude, the averaged map may allow for SIS if,

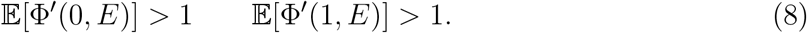

Note that, if the dependence of Φ on *E* and *x* can be separated, Φ(*x, E*) = Ψ(*E*)Θ(*x*), conditions (4) and (7) imply that the system allows for a simple frequency dependent stabilizing mechanism. Assume these conditions are both satisfied, then 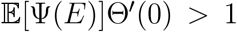 and 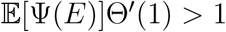 ensure that a system with a fixed fitness 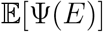 can support stable coexistence, in contradiction to condition 3 (no coexistence under fixed environment) above. Accordingly, stochasticity alone will not induce stability in any system where Φ is linear in *x*_0_ or in *E*; stabilization requires that Φ be a nonlinear and non-separable function. In particular, the logistic map *x*_*t*+1_ = *E_t_x_t_*(1 – *x_t_*) cannot support SIS.

### B. Avoiding diffusive trapping when rare

To understand the conditions for diffusive trapping we again focus on the frequency dynamics when *x* ≪ 1,

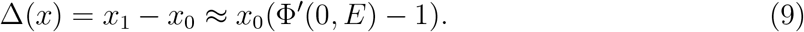

Now Φ′(0, *E*) – 1 is the instantaneous expected growth of the focal species when rare, and at any moment is either positive or negative. For sufficiently small changes in frequency, Δ(*x*), one may use a diffusion approximation [35]; the probability *P* (*x, t*) (of finding the focal species at frequency *x* at time *t*) then satisfies the Fokker-Planck equation (also known as Kolmogorov forwards equation),

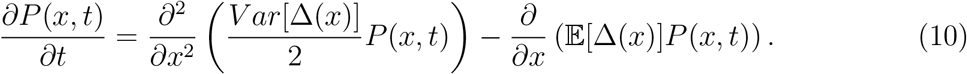

Given (9), when *x* → 0 this equation takes the form

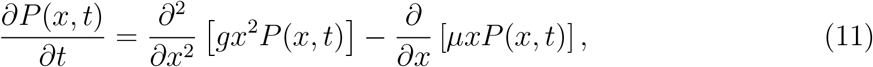

where

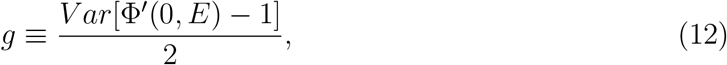

and

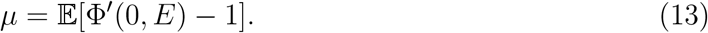

If *P*(*x*) eventually reaches an equilibrium value *P_eq_*, such that *dP_eq_*/*dt* = 0, then as *x* → 0 Eq. (11) implies,

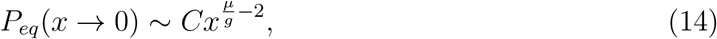

where *C* is a normalization constant. The possibility of diffusive trapping in each extinction zone, regimes *x* ≪ 1 and 1 — *x* ≪ 1, must be considered separately.

Note that we have used only the low-*x* behavior of *Var*[Δ(*x*)] and 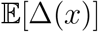, where they depend on the frequency of the focal species alone(we have assumed 1– *x* ≈ 1). To determine the normalization constant *C* one should calculate *P_eq_*(*x*) between zero and one - a more complicated task. Nevertheless, the asymptotic relationship (14) provides the correct **ratio** of the probabilities of finding the system at, say, *x* = 0.001 and *x* = 10^−6^; which is indeed 0.001^(*μ/g*-2)^/10^−6(*μ/g*-2)^.

Coexistence implies that the equilibrium probability distribution can be normalized [47].

Accordingly, coexistence requires

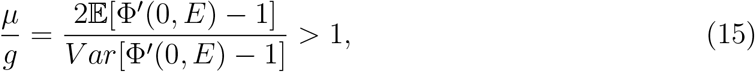

so that *P*(*x*) does not diverge in the extinction zone faster than, or equal to, *x*^−1^. As explained in Appendix I, this binary condition is exactly equivalent to the condition 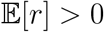 when 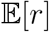 is the mean logarithmic growth rate when rare. Moreover, and unlike 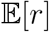 [see Appendix I], when *μ/g* > 1 its numerical value is related to invasion probability (Eq. 49) and also to the persistence time of the system (Eq. S8).

## III. SIS IN DISCRETE TIME MAPS: A SYNTHESIS

In this section we provide four examples of SIS in discrete maps, and show how they all synthesized in a single framework using *μ/g*. For the sake of simplicity we consider a system that jumps between two *E* states (dichotomous noise) but the generalization of our treatment to any type of stochasticity is straightforward.

### A. The lottery model

Chesson & Warner [12] introduced the lottery model. In it, a fraction *δ* < 1 of the population dies and is replaced according to probabilities that reflect the species abundances weighted by their fitnesses. *E* is picked at random at each step, so *δ* as defined here is indeed the correlation time of the environmental variation.

To be more specific, consider a forest with only two tree species in which:

- A fraction *δ* of the adult trees die at each step.
- Each dead adult tree is replaced by a single seedling.
- The ratio between the number of seedlings produced by an individual of the focal species and the number of seedlings produced by an individual of its rival species is *E*.

The chance that the focal species replaces a dead adult tree is

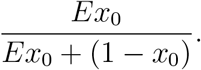

The death toll for the focal species is *δ*_*x*0_. Therefore,

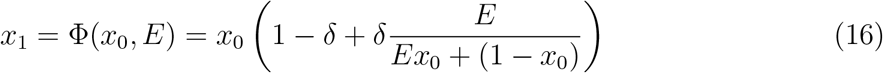

Note that Φ satisfies all the conditions (1-3) above: *x*_1_ = 0 if *x*_0_ = 0, *x*_1_ = 1 if *x*_0_ = 1, *x*_1_ > *x*_0_ if *E* > 1 while if *E* < 1 then *x*_1_ < *x*_0_.

The averaged-map dynamics promote invasion by the focal species when condition (4) is satisfied,

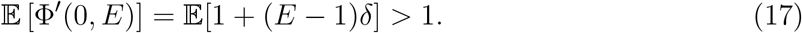

This requires 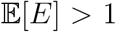, since by definition *δ* > 0. On the other hand condition (7) for the rival species to invade is,

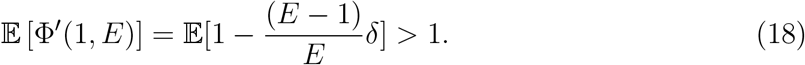

which requires 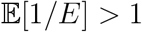.

If *E* = 1/2 with probability 1/2 and *E* = 2 with probability 1/2, then 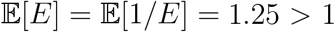 and, with both conditions (4) satisfied, the average map favors SIS. Several more examples with E jumping between 0.2 and 5 and between 0.1 and 10 are illustrated in Figure 1.

Given Figure 1, one might expect SIS to become more and more effective as *δ* grows. The truth is just the opposite: the higher the value of *δ*, the weaker the SIS. To understand this effect we need to consider the impact of diffusive trapping.

Set *E* = *e^s^*, so that *E* = 1 corresponds to *s* = 0, and assume that *s* ≪ 1. Accordingly, Φ′(0, *E*) – 1 = *δ*(*e^s^* – 1) ≈ *δ*(*s* + *s*^2^/2). For a two state system where s flips between +*σ* and –*σ*, the first nonvanishing terms for the expected mean and expected variance of Φ′(0, *E*) – 1 are,

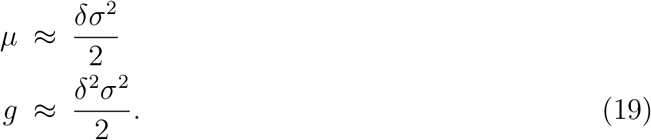

so *μ/g* = 1/*δ*. For small *x*,

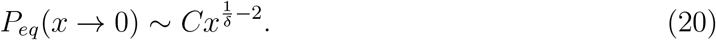

Accordingly, SIS promotes coexistence (*P_eq_* is normalizable) as long as *δ* < 1. The probability of finding the system in the vicinity of the extinction point, which is the effect of diffusive trapping, becomes stronger and stronger as *δ* increases. When *δ* = 1 the probability distribution function can no longer be normalized, meaning that the stabilizing effect has been killed (see Section V). That the expected growth *μ* increases with *δ* does not mean that the stabilizing effect becomes stronger; what is important is the ratio between *μ* and *g*, and this ratio, 1/*δ*, shrinks as *δ* increases, meaning that smaller *δ*s strengthen SIS. In Appendix I we show that 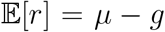. This implies that the condition *μ/g* = 1/*δ* > 1 is identical with the condition 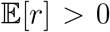, so the threshold for invasion is the same. However, as explained Appendix I, while an increase in *μ/g* implies better persistence properties (see Eq. 49) and a higher probabilities of invasion (Eq. S8), an increase in 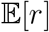 is not necessarily correlated with these properties.

Figure 2 shows that the stabilizing effect weakens as *δ* becomes larger and larger. The equilibrium distribution is concave when *δ* < 1/2 and the system rarely visits the vicinities close to zero and one. On the other hand, the equilibrium distribution is convex with peaks at zero and one when *δ* > 1/2. While these peaks can be normalized if *δ* < 1, they nevertheless imply that the system spends a lot of time close to the extinction zones. Finally at *δ* = 1/2 the value of *P_eq_* is independent of *x* (Fig 2, middle panel, see Danino *et al.* [18]). The concave and convex properties of the distribution have to do with the time to extinction, this also discussed in Section V.

**FIG. 2:**
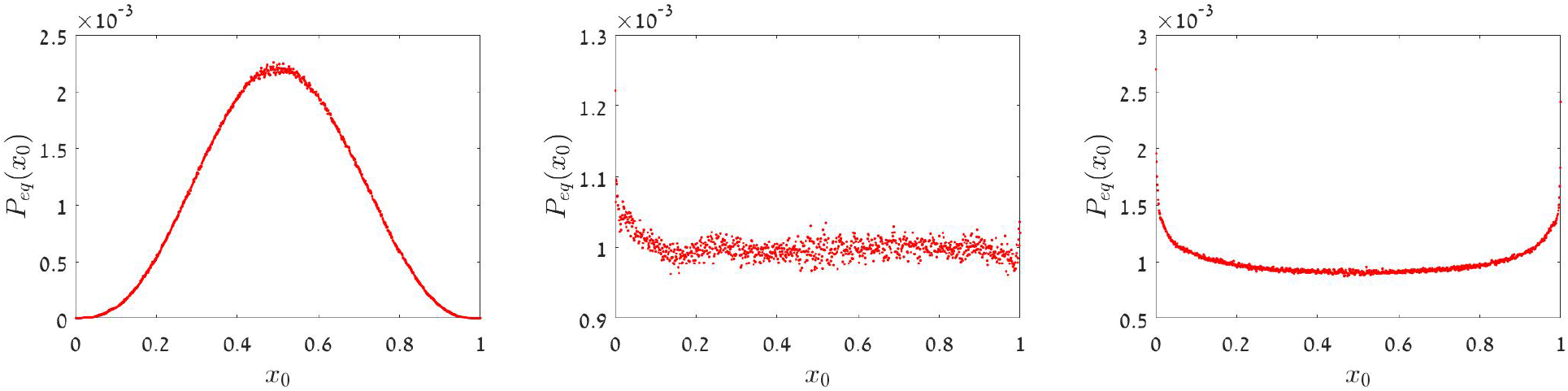
*P_eq_* as a function of *x*, as obtained from numerical simulation of Chesson’s lottery map, *x*_*t*+1_ = *x_t_*[1 – *δ* + *δE*/(*E_x_* + (1 – *x*)], when *δ* = 0.2 (left), *δ* = 0.5 (middle) and *δ* = 0.54 (right). The value of *E* is picked at random in each step to be either *e*^0.3^ or *e*^−0.3^, the process was iterated 10^7^ times, starting from *x* = 1/2. Plotted here are histograms of the number of visits in each bin (bin size 0.001) between zero and one. As predicted, the distribution is concave for *δ* < 0.5, and convex above this value (we have chosen *δ* = 0.54 since for even higher *δ*s the distribution peaks at zero and one are too high and blur the features of the distribution). When *δ* = 0.5 the predicted probability distribution is flat. In that case the outcome of the numerical experiment is, of course, more noisy but still indicates an *x*-independent distribution.

### B. The serial transfer regime

Consider a single microbial species growing exponentially for time *T* with growth rate *r* until the carrying capacity is reached. After this growth phase the population is diluted by a factor *f* into fresh medium and the process reiterated. If *N* denotes the population size immediately after the transfer, then the iterative map is,

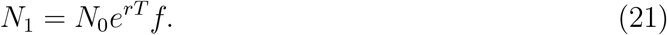

Now consider two competing species. Starting from *N_A_* and *N_a_* = *N*_0_ — *N_A_* individuals, each population grows exponentially (with rates *r_A_* and *r_a_*) until the total population reaches its carrying capacity of *K* individuals. The community is then diluted into fresh medium (dilution factor *f*, where *N*_0_ = *fK*) and regrown. Note that the dilution step never alters the ratio of the competitors, *N_A_*: *N_a_*. Given *N_A_* and *N_a_*, the time *T* until the total population reaches *K* is determined by,

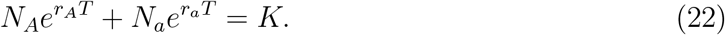

Setting *N_A_* ≡ *x*_0_*N*_0_, *N_a_* ≡ (1 – *x*_0_)*N*_0_, allows *K* to be factored out yielding,

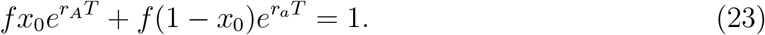

Once *T* is found, the discrete time map for *x* is,

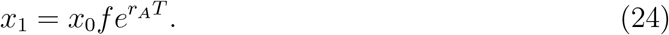

This procedure can be iterated numerically: given *x*_0_, *f*, *r_A_* and *r_a_*, the transcendental equation (23) is solved for *T* which is then plugged into (24) to yield *x*_1_ and so on. Species *A* wins as long as *r_A_* > *r_a_* (*x*_*i*+1_ > *x_i_* for every *x_i_*) and loses as long as *r_A_* < *r_a_* (*x*_*i*+1_ < *x_i_* for every *x_i_*). Moreover if *x*_0_ = 0 then *x*_1_ = 0 and if *x*_0_ = 1 then *x*_1_ = 1 [this follows from Eq. (23)]. The serial transfer regime therefore satisfies conditions (1-3).

We would like to study the dynamics of this system when the values of *r_A_* and *r_a_* fluctuate in time, and in particular when their ratio *R* = *r_A_*/*r_a_* is changed randomly. Although we cannot solve Eq. (23) in general, we can focus on the behavior of *x* when *x* ≪ 1 (i.e., we can study invasion by species *A*). The problem is symmetric so our analysis is relevant to invasion by species *a* as well.

When *x*_0_ ≈ 0 and 1 – *x*_0_ ≈ 1, Eq. (23) implies *e^r_a_T^* = 1/*f* or *T* = – ln(*f*)/*r_a_*. Plugging that into (20) yields,

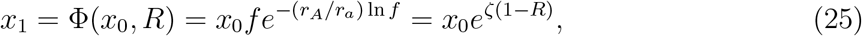

where *R* = *r_A_*/*r_a_* and *ζ* ≡ ln *ζ*. The dilution factor is smaller than one so *ζ* < 0. If *R* is fixed in time and *R* > 1 then *x* invades, while if *R* < 1 then *x* cannot invade. The expected growth of the focal species is positive when *R* varies if,

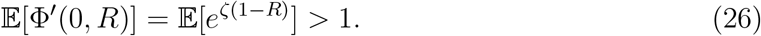

As before, this condition is satisfied if we assume that *R* can take only two values, *R* = exp(±*σ*). As the model is symmetric, fixation is prevented if 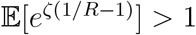. Again, if in half of the cases *R* = exp(*σ*) and in half of the cases *R* = exp(–*σ*), then both conditions (8) hold and the averaged dynamics appears to promote SIS.

To determine the conditions needed to avoid diffusive trapping set *σ* ≪ 1 so that,

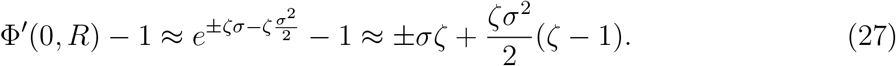

Now,

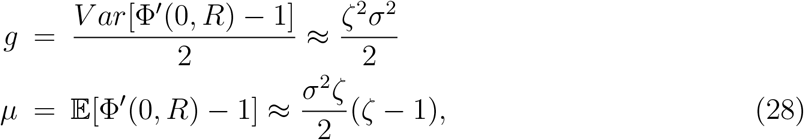

and to the leading order

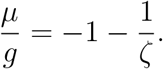

Hence,

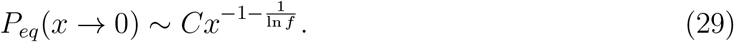

As *f* < 1 so ln *f* < 0, Eq. (29) implies that the two species, A and *a*, coexist **for any level of dilution**. This means that SIS in the serial transfer model promotes coexistence even when the environmental correlation time far exceeds the doubling time. When *f* = 1/*e* ≈ 0.3679 the power is zero and the probability distribution is uniform. For lower dilutions (*f* > 1/*e*)*P_eq_* vanishes close to zero and this produces concave probability distributions. Higher dilutions, for which *P_eq_* diverges close to zero, produce convex distributions. These behaviors are illustrated in Figure 3.

**FIG. 3:**
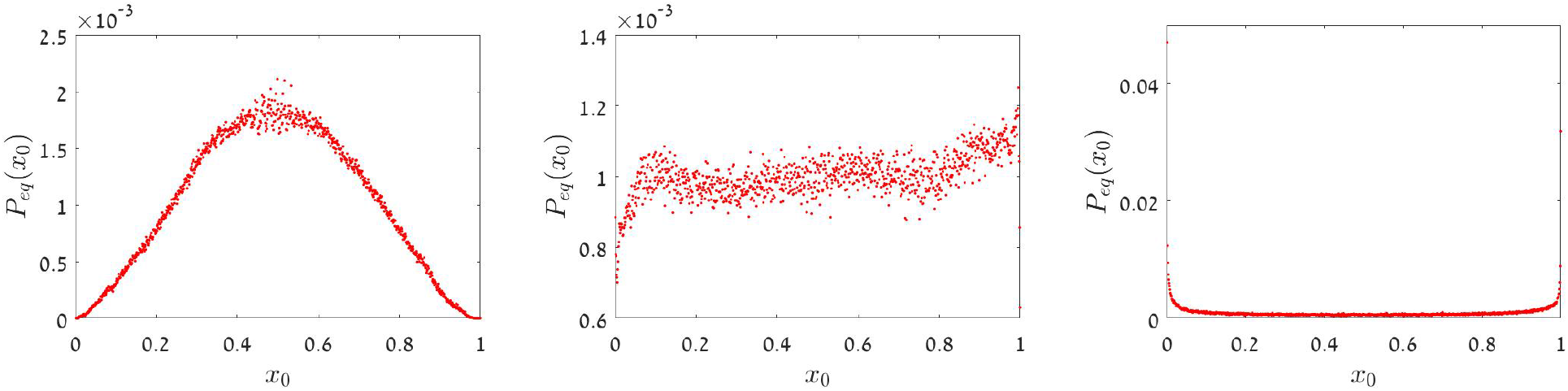
*P_eq_* as a function of *x*, as obtained from numerical simulation of the serial transfer map, for *R* that may take two values, *R* = 1. 1 and *R* = 1 / 1.1. The dilution factors are *f* = 0.7 (left panel) *f* = 1/*e* (middle panel) and *f* = 0.1 (right panel). For each *x*_0_ the value of *T* was determined from Eq. (23) and *x* was incremented according to (21). Starting from *x* = 1/2, the process was iterated 800000 times and the number of visits at each 10^−3^ bin was counted (see caption to Fig. 2). As expected fluctuations are more pronounced in the middle panel, when theory predicts a flat distribution.

### C. Diploids with dominance

[26] studied the dynamics of two alleles, with *A* always dominant to *a*, in a randomly mating diploid population. If the fraction of a alleles in the gamete pool is *x* (and the fraction of *A* is 1 – *x*) then, after random mating, the zygote genotypes follow classic Hardy-Weinberg proportions, with *AA*:*A_a_*: *aa* as (1 – *x*)^2^: 2*x*(1 – *x*): *x*^2^. Set the fitnesses of *AA* and *Aa* to one and the fitness of aa to *W*. Then the frequency of *a* alleles in the next generation’s gamete pool is,

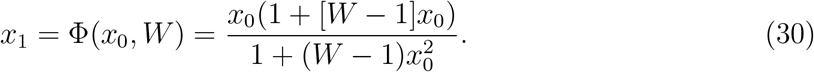

Dominance introduces asymmetry to the model and so invasions by *a* (*x* ≪ 1) and *A* (1 – *x* ≪ 1) must be considered separately. Neither case is trivial.

Invasion by *a* does not quite fit our prescribed framework, which assumes that Φ′(0, *W*) –1 is either positive or negative, because map (30) corresponds to the marginal case with Φ′(0, *W*) = 1 for any *W*. To find the conditions for invasion we need to take the next term in the series,

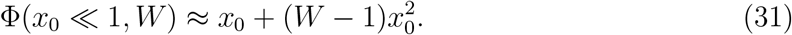

Clearly, invasion by *a* requires 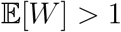.

What about diffusive trapping? Given,

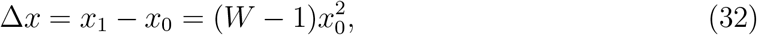

the relevant Kolmogorov forwards diffusion equation at small *x* is,

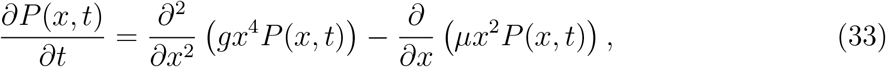

where *g* ≡ *Var*[*W* – 1]/2 and 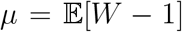. If *W* = exp(±*σ*) and *σ* is small then both *g* ≈ *σ*^2^/2 and *μ* ≈ *σ*^2^/2. The *σ* factors cancel and in the steady state (when the time derivative vanishes) the probability *P_eq_*(*x*) of finding the system at *x* satisfies,

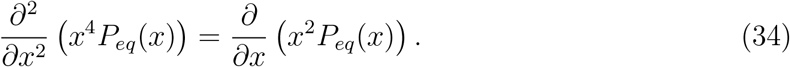

Eq. (34) has to be integrated twice. The constant of the first integration is dropped since it yields a term that diverges like 1/*x*^3^ close to zero. The resulting first order equation is exact and its solution is,

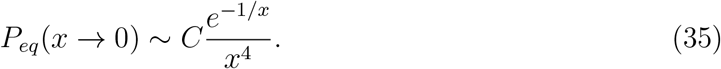

The exponential term vanishes rapidly (faster than any power) as *x* approaches zero and so the system cannot approach the extinction zone near *x* = 0. Allele *a* cannot go extinct.

Allele *a* is a “super persistent” (although slow) invader for which diffusive trapping is strongly prohibited. This happens because only aa homozygotes have fitness *W*, so the growth/decay rate of *a* is a quadratic function of their abundance. When *A* is common the aa homozygotes are exceedingly rare and so, with most genotypes either *AA* homozygotes or *Aa* heterozygotes and having the same fitness of one, selection is inefficient.

On the other hand, when *x* is close to one,

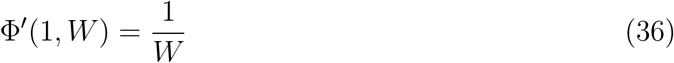

If *W* = exp(±*σ*) then 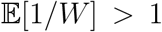, and the expected growth of the *A* allele is positive when rare. However, growth by *A* is linear in (1 – *x*). Once again *μ* = *g* = *σ*^2^/2 and the equilibrium distribution close to *x* = 1 diverges as (1 – *x*)^−1^. The system is trapped: A goes extinct and a fixes [34].

Haldane and Jayakar considered only the expected growth and concluded that both *a* and *A* could invade when rare. Yet on taking diffusive trapping into account one finds that coexistence is not possible in this system; *a* always wins. This conclusion is illustrated in Figure 4.

**FIG. 4:**
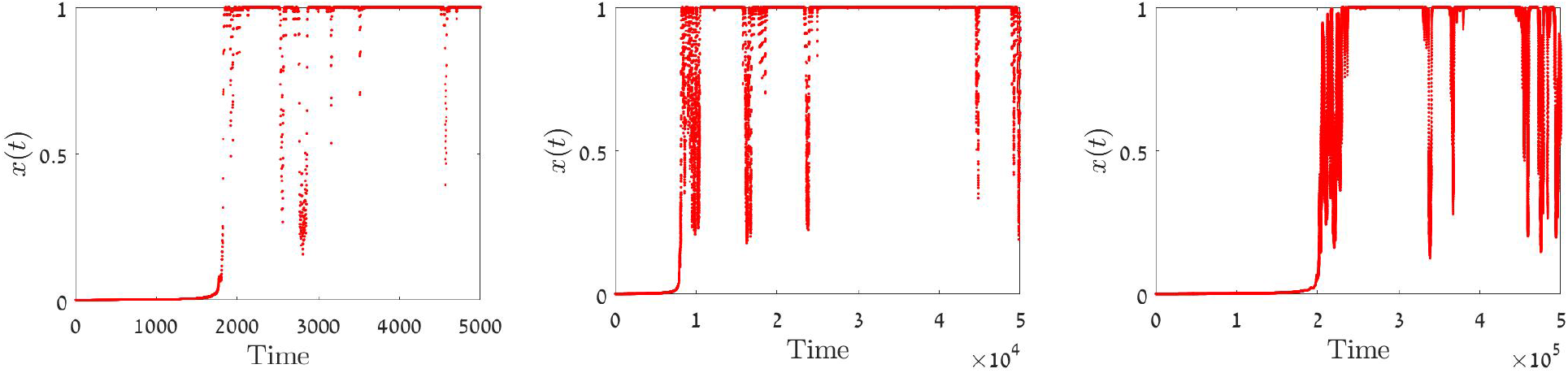
Typical trajectories of *x*(*t*) as a function of t (number of generations) for Haldane-Jayakar map, Eq. (30) with *W* = exp(±*σ*), *σ* = 1 (left) *σ* = 0.5 (middle) and *σ* = 0.1 (right). The initial condition is *x*_0_ = 0.001. Nevertheless, all tra jectories approach *x* = 1 where they remain most of the time (the “bursts” of A allele growth at long times are discussed in Section V). While the expected growth of the rare species is positive at both ends, diffusive trapping nullifies this effect around *x* = 1 and becomes inefficient close to zero.

### D. Continuous time processes

Despite the popularity of Wright-Fisher (nonoverlapping generation) models, generations do overlap. Another class of models, Moran and chemostat models, describe populations with asynchronous births and deaths. In these processes each birth is obligately coupled to a death so that the size of the community, *K* = *N_A_* + *N_a_*, remains fixed. During each elementary time step a species with abundance *N_A_* grows to *N_A_* + 1 (or shrinks to *N_A_* – 1) while its competitor shrinks from *N_a_* to *N_a_* – 1 (or grows to *N_a_* + 1).

We now have three time scales: 1) the time between elementary birth-death events, 2) the community generation time defined as *K* birth-death events and 3) the correlation time of the environment, *δ*, measured in units of a generation (e.g. if *K* = 10^6^ and *δ* = 0.01, the correlation time of the environment is 10, 000 birth-death events).

Now let us specify two microscopic versions of the Moran model. In both of them the log-fitness of a species, *s*(*t*), is a stochastic process with mean *s*_0_ and variance *σ*^2^.

The **local competition** model [43] describes the dynamics of two competing species where a chance encounter between two individuals ends up in a struggle over, say, a piece of food, a mate or a territory. To model that we pick, in each elementary birth-death event, two random individuals for a “duel”, the loser dies and the winner produces a single offspring. If the focal species is represented by *N_A_* individuals, so its fraction is *x* = *N_A_*/*K*, then the chance for an interspecific duel is 2*x*(1 – *x*). The probability the focal species wins a duel (*x* → *x* + 1/*K*) is *p* = 1/2 + *s*(*t*)/4 and the probability it loses (*x* → *x* – 1/*K*) is 1 – *p*. Accordingly, the change in the mean value of *x* in a single birth-death event (1/*K* th of the generation time) is given by,

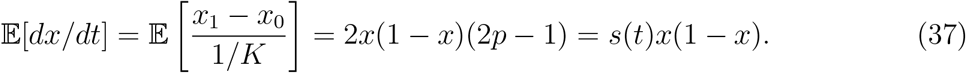

In the **global competition** model [19–21, 43, 44] an individual is chosen at random to die and then all the other individuals compete for the empty slot. The chance the focal species wins is proportional to its relative abundance weighted by its relative fitness; its chance to increase its abundance by one is (1 – *x*)*xe*^*s*(*t*)^/[*xe*^*s*(*t*)^ + (1 – *x*)]. In such a case,

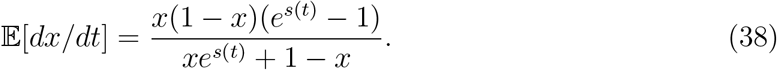

The two models are mathematically equivalent when there is no selection; setting *s*_0_ = *σ*^2^ = 0 defines the same purely demographic (neutral) process 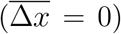 where the focal species has an equal chance, *x*(1 – *x*), to increase or decrease by one individual during each and every elementary birth-death step. If *s* is fixed in time the two models differ slightly, but they both yield the same qualitative results: the focal species grows monotonically to fixation if *s* > 0 and shrinks monotonically to extinction when *s* < 0.

However, the two models behave very differently when the environment fluctuates. To demonstrate that, let us consider the symmetric case: there is no directional selection, *s*(*t*) takes on only two values, ±*σ*, and after each fixed period of time *δ* (after *Kδ* elementary birth-death events) the system picks its state at random, either plus or minus. Since the model is symmetric and we need only analyze invasion by *x*.

For the local competition model, when *x* is small *ẋ* = ±*σx* and so

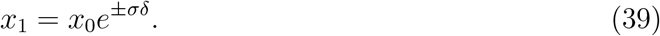

This implies

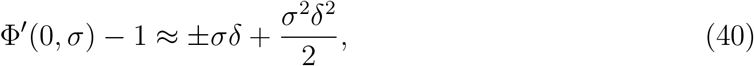

assuming *σδ* ≪ 1. Accordingly, *μ* = *g* = *σ*^2^*δ*^2^/2 and once again diffusive trapping prevents either species invading. Stochasticity does not promote coexistence when competition is local.

In contrast, the global competition model [16, 43] yields,

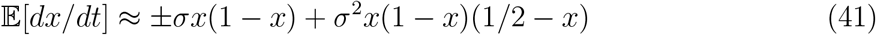

when *σ* ≪ 1. It is the second term that makes the difference. When *x* ≪ 1 the population size at time *δ* is,

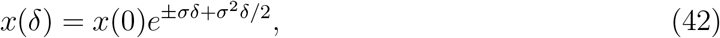

so,

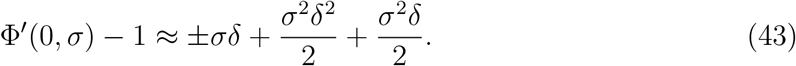

Accordingly,

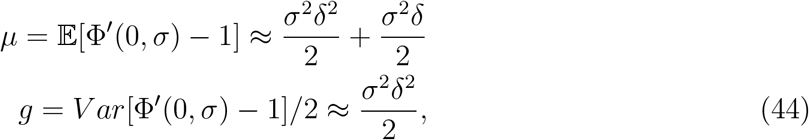

hence

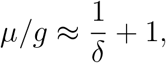

and

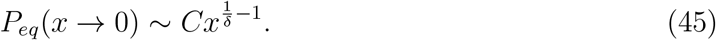

Since *μ/g* > 1, global competition allows the focal species to invade. By symmetry, its rival can also invade from rarity. Stochasticity promotes coexistence when competition is global. Like the serial transfer model, and unlike the lottery model which becomes unstable when *δ* = 1, global competition promotes coexistence for any value of *δ*, even when *δ* is larger than the generation time.

## IV. BET-HEDGING MECHANISMS AND STOCHASTICITY-INDUCED STABILIZATION

In all the cases considered so far, *P_eq_*(*x* → 0) depends only on the correlation time *δ* and not on the amplitude of environmental variations, *σ*, which factors out of the results. This cancellation is attributable to our approximation scheme. We *assumed* that *σ* ≪ 1, and only then calculated the leading contributions to the expected growth *μ* and to its variance *g*. As both quantities scale with *σ*^2^, and as *P_eq_* depends on their ratio, so the dependence on *σ* disappears from the final results. On the other hand the mean is linear in *δ* while the variance scales with *δ*^2^, so *P_eq_*(*x* → 0) depends on *δ*.

However, the rate at which the system approaches equilibrium is strongly affected by *σ*. For example, the forwards Kolmogorov equation for the lottery model, Eq. (11), has the form [18],

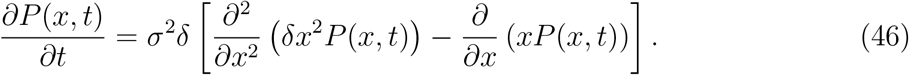

So while *P_eq_* (determined on equating what is written in the squared bracket to zero) is *σ* independent, the factor *σ*^2^*δ* determines the time to equilibrium. The smaller the *σ*, the longer the timescale. Clearly, when *σ* = 0 there is no stochasticity induced stabilization, and so in the absence of environmental variation(*σ* = 0) the system never reaches *P_eq_*.

In practice, when *σ* becomes too small the effect of abundance variations induced by demographic stochasticity can no longer be neglected, no matter how large *K* is [18]. When demographic fluctuations dominate SIS, stability is lost. The relationships between *σ*, invasibility and persistence time are more complicated when demographic stochasticity is important.

The effects of decreases in *δ* are less obvious. On the one hand they slow convergence (the term *σ*^2^*δ* in Eq. (46) becomes smaller). Yet on the other the *x*^1/*δ*^ behavior of *P_eq_* implies that the smaller the *δ*, the smaller the chance of finding the system in the extinction zone. In most cases (as discussed in the next section), decreases in *δ* strengthen SIS.

These insights help explain the relationship between SIS and bet-hedging mechanisms. Since the original lottery model works only when generations do not overlap, many authors suggested a general insight, namely that diversity is promoted by mechanisms “for persisting during unfavourable periods, such as a seedbank, quiescence or diapause.” [2]. The idea is that long-lived life-history stages that shield individuals from competition promote diversity by buffering “species against sudden rapid declines” [42]. As we have seen [and as was already pointed out by Abrams [1]] overlapping generations are not necessary for SIS. What can be said about bet-hedging strategies in general?

Bet-hedging mechanisms ar act to buffer growth rates against environmental variations. For example, one may compare two species that satisfy *N*_*t*+1_ = *λ_t_N_t_*, where *λ* is the yearly growth factor. Without bet-hedging the *λ* of a certain species may be 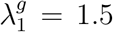 in good years and 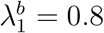 in bad years, while a successful bet-hedging strategy implemented by its competitor yields a fixed growth rate of *λ*_2_ = 1.1 in all years. Each year is either good or bad with probability 1/2.

Although the expected arithmetic growth of the bet-hedger is lower than that of its rival 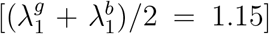, its geometric mean growth rate is higher 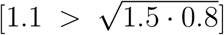, and this implies that the bet-hedger population is larger (with probability one) in the long run [40]. Accordingly, in the absence of density-dependent population regulation the bet-hedger population wins the competition.

In the models considered above density dependence plays an important role, and in most of them the two species are playing a zero-sum game. Therefore, growth rate fluctuations of one species are exactly mirrored by the growth rate fluctuations of its rival. In such a case bet-hedging strategies that buffer seedling recruitment or egg production against environmental variations lead to a decrease in the effective strength of environmental stochasticity, i.e., to smaller *σ*^2^. This by itself *weakens* SIS.

Nevertheless, bet-hedging *may* bring about SIS. Many of the bet-hedging mechanisms, like seed banks or adult longevity, lead to an increase in the effective generation time. Since *δ* is the ratio between the correlation time of the environment and the generation time, its value decreases. As we have seen any decrease of *δ* contributes substantially to the efficiency of SIS, so this side effect of bet hedging may increase a system’s stability.

## V. PERSISTENCE TIME AND EXTINCTION DYNAMICS

Until now we have classified systems into two groups, those that support stochasticity induced stability and those that do not. Our criterion was based on the behaviour of *P_eq_*(*x*) close to the extinction points *x* = 0 and *x* = 1. If *μ*/*g* > 1 then *P_eq_*(*x*) ~ *x*^*μ*/*g*−2^ (or a similar expression close to *x* = 1) is normalizable and the system supports SIS. Otherwise *μ*/*g* ≤ 1 and *P_eq_* diverges in the extinction zone so fast that the equilibrium probability distribution cannot be normalized and so the system is unstable.

But what is the practical meaning of all this? At the end of the day, biodiversity and polymorphism reflect the balance between the rates of extinction and speciation/mutation. What is the relevance, then, of the properties of *P_eq_* to the rates of loss of species and alleles? One might expect that extinction rates should increase given strong support for *P_eq_* in the “danger zones” close to fixation and extinction. Still, the question remains how to quantify this phenomenon. Another trivial insight is that the chance of extinction for a single individual (a mutant or an invader, say) is larger than the chance of extinction when *x* = 1/2, say. Again one would like to quantify the effect of the initial conditions on extinction rates.

We need to consider finite systems if we are to quantify extinction rates. In the last sections we treated *x* as a continuous variable for an infinite community, but no species can actually go extinct in an infinite community. Only by recognizing that individuals are discrete, and that communities are finite, can our intuition about stability and extinction rates be translated into quantitative, or at least semi-quantitative, predictions.

For a finite community with *K* individuals, the frequency of the focal species, *x*, can only change by units of ±1/*K*. This property manifests itself in *demographic stochasticity*, the endogenous abundance fluctuations that reflect the stochasticity inherent to the underlying birth-death process [38]. We will ignore all the effects of demographic stochasticity save one, namely that at any given time *t* the rate of extinction is proportional to the chance that the system visits the extinction zone *x* < 1/*K* [11, 37, 43].

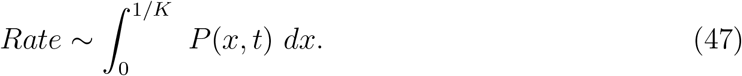

Moreover, we will assume that *K* is sufficiently large that factors not scaling with *K* can be neglected.

For a **stable** system *P_eq_* exists, and after some period of time the probability distribution converges on *P_eq_*. With *K* large, the rate of extinction given by Eq. (47) is very small; the system “leaks” to extinction through a tiny “hole” between zero and 1/*K*. Qualitatively, if *P_eq_* ~ *Cx*^*μ*/*g*−2^ and *μ*/*g* > 1 then the rate of extinction is proportional to,

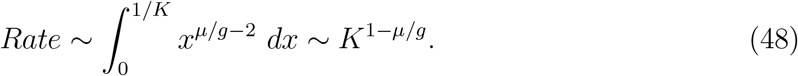

This is another aspect of stability; when *μ*/*g* > 1, the rate of extinction decays to zero as *K* goes to infinity. The mean time to extinction, *T*, is inversely proportional to the rate of extinction and so the *K* dependency of *T* is given by,

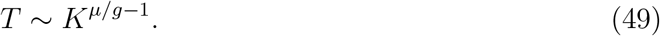

For the models described so far the time to extinction behaves like a power-law in *K* (see discussion). In particular it has been shown for the Moran process considered above [16, 29] that

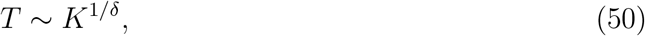

in agreement with Eq. (45).

Of course the system may reach extinction before it relaxes to *P_eq_*. However, the large *K* dependency of *T* is not affected by the initial conditions in the systems we have considered [16, 43]. There may be losses during the relaxation process, but these do not scale with *K* and so they only change result (49) by a constant factor. This implies that the chance that a single mutant becomes established, which is related to the difference between *T*(*N*_0_ = 1) and *T*(*N*_0_ ≫ 1), is also *K*-independent, as has been proved in [43]. See Appendix I (Eq. S8) for a more qualitative analysis of the chance of invasion.

A different route to calculating persistence times must be used for **unstable** systems that lack SIS. With *P_eq_* no longer normalizable the system never equilibrates and so the equilibrium probability distribution *P_eq_* does not exist. To understand the extinction dynamics in such a system consider the simple, famous and solvable model,

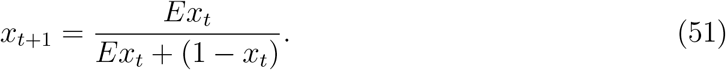

As we have seen, this is the lottery model in the limit *δ* = 1.

The quantity *u* ≡ (1 − *x*)/*x* satisfies *u*_*t*+1_ = *u_t_*/*E*, and so *z* = ln *u* obeys the recurrence relation

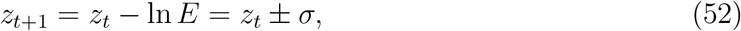

where the last identity holds for telegraphic noise with *E* = exp(±*σ*).

Accordingly, the variable *z* performs a simple random walk, between −∞ and ∞, with step size *σ*. The solution in the continuum limit is given by the diffusion equation with diffusion constant *σ*^2^. If the initial condition is *x*(*t* = 0) = 1/2, then *u*(*t* = 0) = 1 and *z*(*t* = 0) = 0, so the probability finding the system at *z* at time *t* is given by,

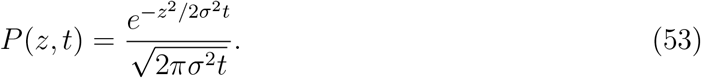

Since *x* = 1/(1 + *u*) = 1/(1 + *e^z^*), one finds [25],

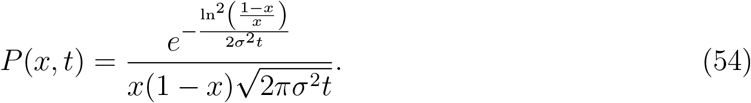

Eq. (54) allows us to understand the extinction dynamics when SIS is inefficient. *P*(*x, t*) is always a normalized and strictly legitimate probability distribution function. However, with the passage of time *P*(*x, t*) develops higher and higher “wings” in the extinction zones as it attempts to to reach its unattainable target, the non-normalizable function *P_eq_* = 1/*x*(1 − *x*) (Figure 5). Technically speaking, the non-normalizable *P_eq_* is the *infinite density* associated with (54) [3]. For each fixed *t* there is a very tiny and ever-shrinking zone, *x* < exp(−*σt*/2), where the probability drops to zero. Diffusion has trapped the system in the extinction zone. As time progresses *x* is found, on average, closer and closer to zero (or to one) until at some stage it crosses the single individual limit at *x* = 1/*K*, where the dynamics halts.

**FIG. 5:**
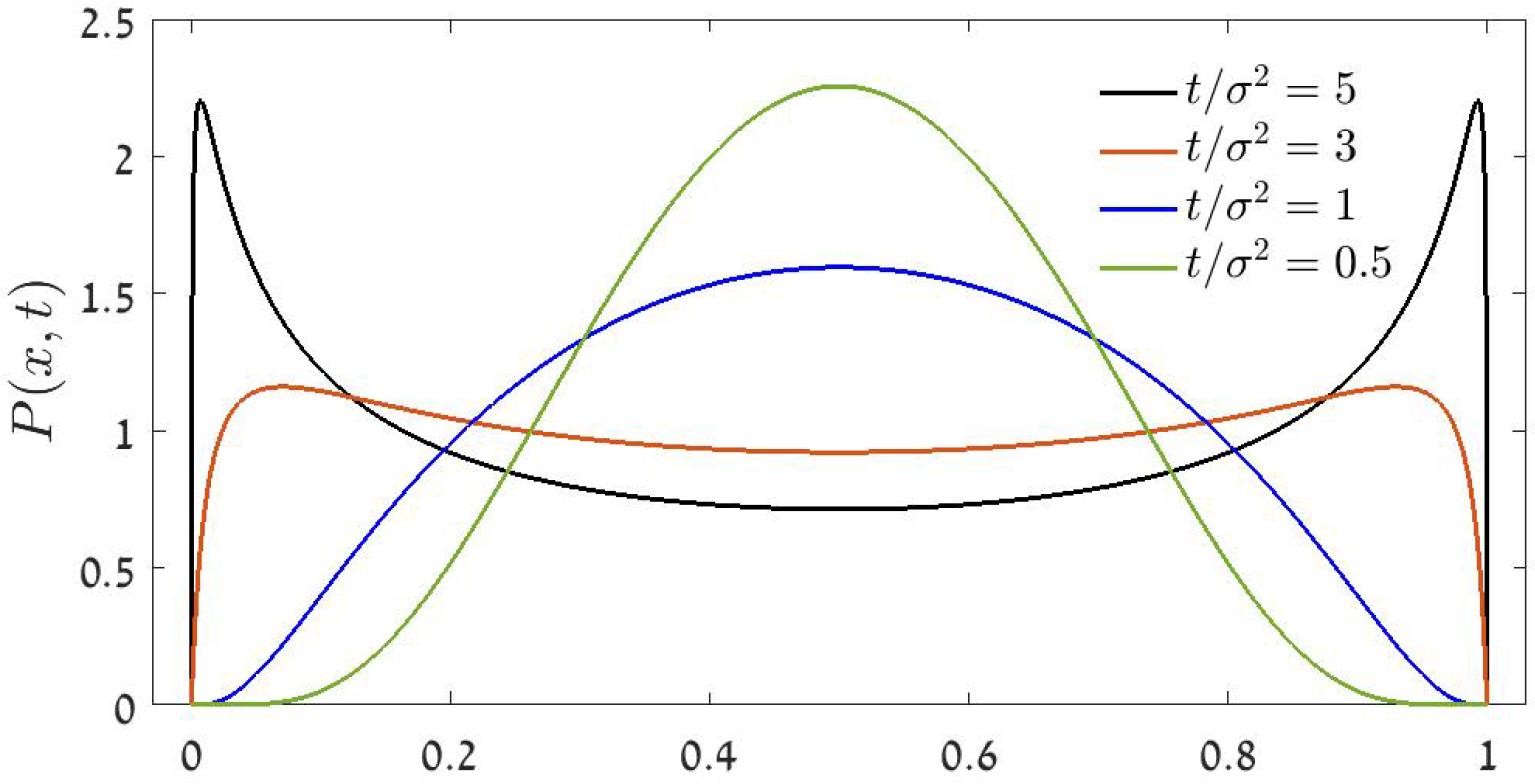
The evolution of the probability distribution *P*(*x,t*), Eq. (54), through time. As *t*/*σ*^2^ grows, the pdf develops wings at the extinction zone, still it drops sharply to zero when *x* or 1 − *x* are smaller than exp(−*σ*^2^*t*/2).

A rare sequence of good years may cause a “burst” of growth even if a population is extinction prone. This phenomenon is clearly seen for the *A* allele in Figure 4. With time the fraction of *A*-s in a typical system decrease and the corresponding probability distribution function (that reflects an average over many specific histories) accumulates closer and closer to *x* = 1. Still, in any specific system, as long as *A* exists, a rare sequence of successive good years may lead to an exponential increase of the *A* population, causing the “bursts” seen in Figure 4.

With time, the *A* population decreases so longer sequences of good years are required to produce a burst. Accordingly, bursts become rarer and rarer and also lower in amplitude until the very last individual dies. On the other hand, after a successful “rejuvenative burst” the chance for another burst increases and so bursts tend to cluster. Accordingly, a typical abundance time-series of an extinction prone allele is highly nontrivial, with an intricate burst structure that may provide false clues to long-term dynamics.

## VI. SIS IN HIGH-DIVERSITY ASSEMBLAGES

SIS weakens as the number of species is increased beyond two. Let us present a few results and considerations that have to do with this insight.

Chesson and Hatfield [28] considered the lottery model when the number of species is *S*. Again, a fraction *δ* of the population is killed and replaced by the offspring of a species where abundance is weighted by fitness, so

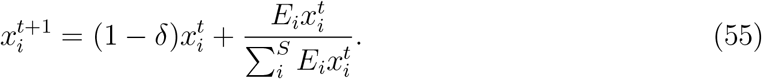

When this model is symmetric, or “time-averaged neutral” (*E_i_*-s are picked from the same distribution for all species *i* = 1..*S*), the equilibrium multivariate distribution *P_eq_*(*x*_1_, *x*_2_..*x_S_*) was found [28]. The corresponding single species distribution *P_eq_*(*x*|*S*), obtained from the multivariate function by integrating out *S*−1 of the *S* species, was calculated in Danino *et al*. [18]. When sufficiently strong, SIS causes *P_eq_*(*x*|*S*) to be sharply peaked around *x* = 1/*S*. This is a simple consequence of symmetry in the model; each species has the same average fitness and so the abundance of each must fluctuate around 1/*S*. As *x* → 0 the *x*-dependence of *P_eq_*(*x* → 0) switches from its two species form, *Cx*^−2+1 /*δ*^ (Eq. 20), to

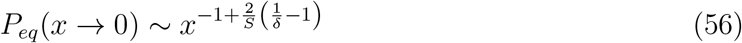

and the time to extinction of each species is shortened significantly.

In an open system new types emerge through speciation, mutation or migration from a regional pool. This leads to an increase in the number of species and as a result SIS becomes weaker and the rate of extinction increases until the system balances. This phenomenon was discussed recently, both numerically and analytically [17, 18, 21, 31] using the continuous time Moran models presented above. When all parameters (like *σ* and *δ*) are equal, the species richness in a global competition process is higher than in a local competition process due to SIS. However, the condition for positive response of biodiversity to stochasticity (an increase in richness/polymorphism with *σ*) is *δ* < 2/Θ, when Θ is the fundamental biodiversity parameter *νK* and *ν* is the rate at which new species emerge in the system (e.g., the mutation rate).

## VII. ASYMMETRIC COMPETITION

So far we have considered only the symmetric case, where all species have the same time-averaged fitness. In the asymmetric case the competitor with the higher mean fitness is more likely to reach fixation. Nevertheless, the system may still support SIS if the mean fitness difference is sufficiently small [16, 20, 21, 43].

The fixation/extinction times for two species competition in a fixed environment are logarithmic in *K*. When *E* is time-independent, the time needed for a single mutant with *E*=*e*^*s*_0_^ to reach fixation against a species with *E*=1 is *T_F_* ~ln *K*/|*s*_0_|. On the other hand, when the fitness of a beneficial species is *E* = *e*^*s*_0_±*σ*^ and the deleterious fitness is *E* = *e*^±*σ*^ (so the mean difference in log-fitness is still *s*_0_, but the relative fitness fluctuates) the time needed for purifying selection scales with *K* as

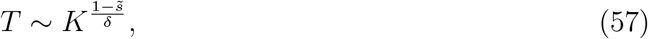

where 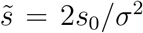 [16, 43]. Eq. (57) holds as long as 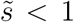, i.e., when the amplitude of environmental fluctuations is large with respect to the mean fitness difference. If 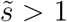 the mean map has no attractive fixed point and there is no SIS.

When competition is asymmetric the chance that the inferior competitor reaches fixation is vanishingly small and the chance of the beneficial species to win is close to one. Nevertheless, both species can invade and the SIS persistence times (given in (57) - note that *T* is measured in generations) may be huge [43].

## VIII. DISCUSSION

Throughout this paper we have tried to provide a concise and intuitive introduction to the phenomenon of stochasticity-induced stabilization. We focused our attention on the simplest cases, most of them involving two species zero-sum competition dynamics with dichotomous (two-state) environmental fluctuations.

We provided a new perspective to the stabilization problem by focusing on the arithmetic mean of the growth when rare, *μ*. Invasion is not guaranteed by the condition *μ* > 0, because *μ* has to be large enough to overcome the effect of diffusive trapping, hence the invasion condition is *μ*/*g* > 1. If the low-frequency dynamics is linear (i.e., if when *x* ≪ 1 one finds *dx*/*dt* ≈ *E*(*t*)*x*) this condition is equivalent to the traditionally used criteria for the mean logarithmic growth rate when rare, 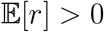, as explained in Appendix I.

There are two advantages to our method: first it allows us to consider cases of nonlinear low-frequency dynamics, like that of a recessive allele where *dx*/*dt* ≈ *E*(*t*)*x*^2^. Moreover, unlike 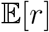 the parameter *μ*/*g* provides a quantitative measure of the invasion probability (Eq. S8) and the persistence time (Eq. 49).

The simplest and most familiar map in population genetics, Eq. (2), which is also the lottery model map with *δ* = 1, is exactly marginal. The stabilizing effect of the expected growth is balanced precisely by the effect of diffusive trapping, and the net outcome [as seen in Eq. (52)] is an unbiased random walk along the ln *z* axis. This marginality manifests itself in the *x*^−1^ divergence of *P_eq_* as *x* → 0 at the threshold between normalizable and non-normalizable distributions.

Stability requires some mechanism to break the tie between growth and trapping. In the lottery model the expected growth scales with *δ* while the variance scales with *δ*^2^, and so growth is stronger than trapping whenever *δ* < 1. The stabilizing effects are even stronger in the corresponding continuous time (Moran) dynamics. Now the asymmetry between growth and decline is larger because the overall demographic benefit to a growing population includes not only births that originated from individuals present at the beginning of the growth period, but also those from individuals born during the period [20, 36].

The marginal process (2) also describes a serial transfer experiment in which dilution always occurs after a fixed time *T* when the populations are still growing. In this case, and as demonstrated experimentally [51], there is no SIS. However if the dilution takes place after the community has reached carrying capacity then, as described in subsection III B above and as demonstrated experimentally [51], SIS promotes coexistence. The reason is that, while the chance that the rare species picks a relatively good or a relatively bad environment is equal, the net time for growth in the good environment is longer than in the bad environment and so the random walk along the abundance axis is biased towards growth.

The diploid model of Haldane and Jayakar represents a hybrid case. The dynamics of A when rare is very similar to Eq. (2). As a result the expected growth and the diffusive trapping are precisely balanced and so A cannot invade. On the other hand the growth and decay of the recessive allele *a* (when rare) are not exponential; they depend on the square of its density. In this case diffusive trapping is relatively weak and invasion is slow but persistent.

Another important insight is the distinction between bet-hedging mechanisms and mechanisms that yield SIS. The simplest outcome of any bet-hedging strategy is, as is implied by the name, to lower the effective amplitude of the environmental stochasticity, i.e. to decrease *σ*. In that sense, bet-hedging strategies act against the stabilizing effect: there can be no stochasticity-induced stabilization without stochasticity. However, in many cases (e.g. seed banks, adult longevity) bet-hedging strategies increase the effective generation time. Essentially, they decrease *δ* (the correlation time of the environment measured in units of a generation), thereby reducing the chances of finding the system in the extinction zone, and so strengthening SIS.

Finally we would like to discuss the qualitative features of the “coexistence” that the system achieves via SIS mechanisms. The word “coexistence” allows for different interpretations. We identified coexistence with the existence of *P_eq_* in the two (or many) species system. A stronger condition, proven by Chesson [7, 27] for the lottery model, is that population fluctuations converge on a stationary stochastic process with all densities positive, i.e., that the system forgets its initial conditions and all the correlation functions eventually reach equilibrium. These features are irrelevant to the stability metric used above, so we did not consider them here.

If *P_eq_* exists, and if the chance of extinction during the transient time (from a generic initial condition until the system equilibrates) is negligible, then the typical persistence time [see Section V and [16, 29]] grows as the power of the community size *K*. In sharp contrast, when the system supports a genuine attractive equilibrium point (cases with density or frequency dependence, or with niche differentiation) the time to extinction grows *exponentially* with *K* [38].

One can infer the power-law behavior from a simple argument [43, 50]. When the system is subject to environmental stochasticity the growth rate of the focal species may become negative, say −*σ*. During long periods of negative growth abundance decays exponentially, *x*(*t*) = *x*_0_ exp(−*σt*). If extinction is declared when *x* = 1/*K*, the time to extinction will be *T_d_* = ln(*Kx*_0_)/*σ*. This time is relatively short, so the bottleneck that determines the persistence time of the system is the probability of picking a period *T_d_* during which the growth rate is always negative. If the growth rate is picked at random every *δ* generations, then this probability scales as exp(−*T_d_*/*δ*) ~ *K*^−1/*σδ*^ and the time to extinction, which is the inverse of this probbaility, grows as a power of *K*.

This qualitative argument is quite powerful, still it relies on two assumptions. One is that the population decline during bad years is exponential. The *a* dynamics in Haldane-Jayakar model violates this assumption because at low densities *dx*/*dt* ~ (*W* − 1)*x*^2^. When *W* is less than one (bad years) the decline to *x* = 1/*K* takes approximately *Kx*_0_ generations (and not the logarithm of this quantity as before). The probability of picking such a sequence of bad years is exponentially small in *K*/*δ*, so the time to extinction will grow exponentially with *K*. This is consistent with Eqs. (35) and (47) above.

Second, the probability of picking a sequence of bad years that leads to extinction with linear growth/decline is exp(−*T_d_*/*δ*). This probability is exponentially small in *K* whenever *δ* scales as *c*_1_/*K* where *c*_1_ is some constant. This has to do with our previous observation that SIS becomes stronger and stronger as *δ* shrinks. A detailed analysis of the time to extinction in that case is presented in [16]. Although these short-term fitness fluctuations, with a correlation time of a few birth-death events, are much smaller in amplitude than the fluctuations that persist for half or tenth of a generation, their effects on stability may be substantial.

## IX. ACKNOWLEDGMENTS

N.M.S. acknowledges helpful discussions with Eli Barkai, David Kessler, Jayant Pande and Niv de-Malach. This research was supported by the ISF-NRF Singapore joint research program (grant number 2669/17).

## Supplementary information

### I. 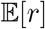 VS. *μ*/*g*: A COMPARISON

In this appendix we will clarify the relationships between our *μ*/*g* analysis and the usage of the mean growth rate when rare, 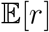, as a measure for invasibility and persistence. When a concrete example is needed we consider the lottery model as defined in Eq. (16).

#### A. Growth rate when rare

First we recite the definition of 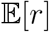, which is the mean growth rate along the log abundance (or log-frequency) axis. If for a certain species the outcome of two successive frequency measurement are *x_t_* and *x*_*t*+Δ*t*_, then by definition,

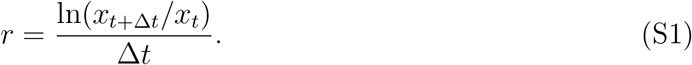

Accordingly, given a time series of (low) frequencies {*x_t_*, *x*_*t*+Δ*t*_, *x*_*t*+2Δ*t*_…} and so on, this mean growth rate is defined as [10],

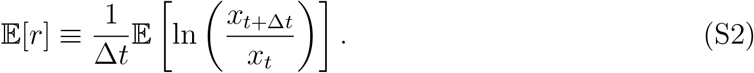

For the discrete time lottery model Eq. (16) above (for *x*_0_ ≪ 1) implies:

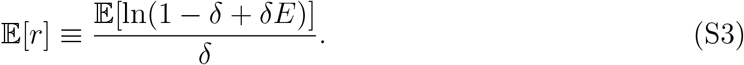

When *E* = *e*^±*σ*^ and the diffusion approximation holds,

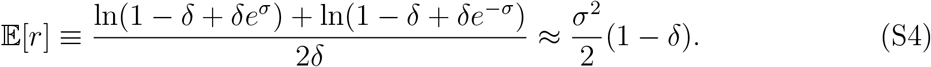

If 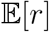 is indeed a quantitative measure of persistence one should expect that persistence time grows and the chance of invasion increases with *σ*. As we shall see, this is not the case.

#### B. Fokker-Planck equation in log-abundance coordinates

Let us consider the Fokker-Planck equation for rare species, Eq. (11),

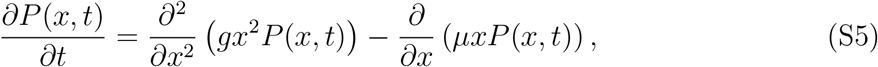

To clarify the relationships between the stochastic process and 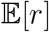, we would like to switch to the log-frequency domain. To do that we substitute

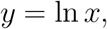

and use *Q*(*y, t*) for the probability distribution function in terms of *y*. Since *P*(*x*)*dx* = *Q*(*y*)*dy* one has,

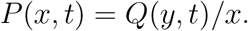

We also switch to measure time in units of a single generation (such that *Q*(*y,t* + *δ*) ≈ *Q*(*y,t*) + *δ∂Q*(*y, t*)/*∂t*). Pugging all that into (S5) one finds,

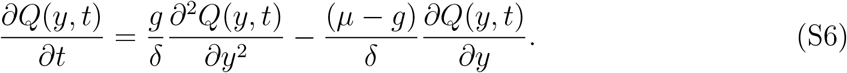

For the lottery model we discovered above (Eq. 19) that *μ* = *δσ*^2^/2 and *g* = *δ*^2^*σ*^2^/2. A comparison with Eq. (S4) yields, for the lottery model, 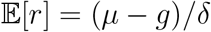, so in that case,

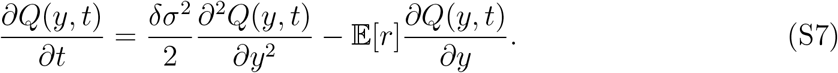

#### C. The chance of invasion

Eqs. (S6) and (S7) are standard convection-diffusion equation, and many results are known in that case. For example it is known [45, Eq. 2.3.8] that if a particle starts at a distance *y*_0_ to the right of the extinction point and it invades only if it reached *y*\* before reaching zero, the chance of invasion 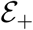 is,

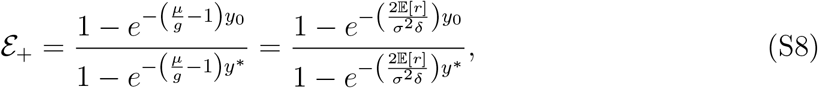

where the right expression holds for the lottery model. In an invasion experiment, if the initial condition is *x* = 2*ϵ* and invasion is declared when *x* reaches *x_f_* before reaching *ϵ*, then *y*_0_ = ln 2 and *y*\* = ln(*x_f_*/*ϵ*).

Accordingly, one observes that the parameter we have used through this paper, *μ*/*g* − 1, is indeed the parameter that sets the chance of invasion from rarity. 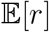, on the other hand, sets the value of the “convection” term (the first derivative, that determines the mean velocity along the logarithmic axis) in Eqs. (S6) and (S7), but its value is not related directly to the chance of invasion since it does not take into account the diffusion (second derivative) term. Only the **ratio** between convection and diffusion, *μ*/*g* − 1, determines the chance of invasion.

In particular, for the lottery model in this parameter regime the actual chance of invasion is independent of *σ*. When *σ* increases while all other parameters are kept fixed, the ratio *μ*/*g* does not change. Although an increase in *σ* leads to an increase in 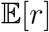, this has nothing to do with invasibility.

A similar problem appears when the persistence time of a system is analyzed. As shown in section V, this time scales like *K*^*μ*/*g*^, so when the diffusion approximation holds it is independent of *σ*, although the numerical value of 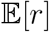 increases with *σ*.

Note that *μ*/*g* and 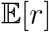 must have the same sign - either both are negative or both are positive. When these parameters are negative the chance of invasion goes to zero with *ϵ* (a natural choice for *ϵ* is 1/*K* [11], so the chance goes to zero when *K* diverges) and the persistence time is *K* independent. On the other hand when these parameters are positive there is still a chance to invade when *K* diverges, and after invasion the persistence time is a positive power of *K*. Accordingly, the change of sign of 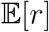 (as well as *μ*/*g*) is a good marker of the transition between these two types of behavior. However, while the sign of 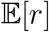 is a good binary indicator, its magnitude does not necessarily provide useful information about the persistence of the community, while the magnitude of *μ*/*g* does.

## References

[1] Abrams, P. (1984). Variability in resource consumption rates and the coexistence of competing species. Theoretical Population Biology, 25, 106–124.

[2] Adler, P.B. (2014). Testing the storage effect with long term observational data. In: Temporal dynamics and ecological process (eds. Kelly, C.K., Bowler, M.G. & Fox, G.A.). Cambridge University Press, p. 82.

[3] Aghion, E., Kessler, D.A. & Barkai, E. (2019). From non-normalizable boltzmann-gibbs statistics to infinite-ergodic theory. Physical Review Letters, 122, 010601.

[4] Bell, G. (2010). Fluctuating selection: the perpetual renewal of adaptation in variable environments. Philosophical Transactions of the Royal Society of London B: Biological Sciences, 365, 87–97.

[5] Caceres, C.E. (1997). Temporal variation, dormancy, and coexistence: a field test of the storage effect. Proceedings of the National Academy of Sciences, 94, 9171–9175.

[6] Cain, A.J., Cook, L. & Currey, J.D. (1990). Population size and morph frequency in a longterm study of cepaea nemoralis. Proc. R. Soc. Lond. B, 240, 231–250.

[7] Chesson, P. (1982). The storage effect in stochastic competition models. Mathematical ecology: Proceedings, Trieste, pp. 76–89.

[8] Chesson, P. (1994). Multispecies competition in variable environments. Theoretical Population Biology, 45, 227–276.

[9] Chesson, P. (2000). Mechanisms ofmaintenance ofspecies diversity. Annual review of Ecology and Systematics, 31, 343–366.

[10] Chesson, P. (2003). Quantifying and testing coexistence mechanisms arising from recruitment fluctuations. Theoretical population biology, 64, 345–357.

[11] Chesson, P.L. (1982). The stabilizing effect ofa random environment. Journal of Mathematical Biology, 15, 1–36.

[12] Chesson, P.L. & Warner, R.R. (1981). Environmental variability promotes coexistence in lottery competitive systems. American Naturalist, pp. 923–943.

[13] Chisholm, R.A., Condit, R., Rahman, K.A., Baker, P.J., Bunyavejchewin, S., Chen, Y.Y., Chuyong, G., Dattaraja, H., Davies, S., Ewango, C.E. et al. (2014). Temporal variability of forest communities: empirical estimates of population change in 4000 tree species. Ecology letters, 17, 855–865.

[14] Cook, L. & Jones, D. (1996). The medionigra gene in the moth panaxia dominulcr: the case for selection. Phil. Trans. R. Soc. Lond. B, 351, 1623–1634.

[15] Crow, J.F., Kimura, M. et al. (1970). An introduction to population genetics theory. An introduction to population genetics theory.

[16] Danino, M., Kessler, D.A. & Shnerb, N.M. (2018). Stability of two-species communities: drift, environmental stochasticity, storage effect and selection. Theoretical Population Biology, 119, 57–71.

[17] Danino, M. & Shnerb, N.M. (2018). Theory of time-averaged neutral dynamics with environmental stochasticity. Physical Review E, 97, 042406.

[18] Danino, M., Shnerb, N.M., Azaele, S., Kunin, W.E. & Kessler, D.A. (2016). The effect of environmental stochasticity on species richness in neutral communities. Journal of theoretical biology, 409, 155–164.

[19] Dean, A.M. (2005). Protecting haploid polymorphisms in temporally variable environments. Genetics, 169, 1147–1156.

[20] Dean, A.M. (2018). Haploids, polymorphisms and fluctuating selection. Theoretical population biology.

[21] Dean, A.M., Lehman, C. & Yi, X. (2017). Fluctuating selection in the moran. Genetics, pp. genetics–116.

[22] Dobzhansky, T. (1943). Genetics of natural populations ix. temporal changes in the composition of populations of drosophila pseudoobscura. Genetics, 28, 162.

[23] Ellner, S.P., Snyder, R.E. & Adler, P.B. (2016). How to quantify the temporal storage effect using simulations instead of math. Ecology letters, 19, 1333–1342.

[24] Fisher, R.A. & Ford, E.B. (1947). The spread of a gene in natural conditions in a colony of the moth panaxia dominula l. Heredity, 1, 143.

[25] Gillespie, J.H. (1972). The effects of stochastic environments on allele frequencies in natural populations. Theoretical population biology, 3, 241–248.

[26] Haldane, J.B.S. & Jayakar, S.D. (1963). Polymorphism due to selection of varying direction. Journal of Genetics, 58, 237–242.

[27] Hatfield, J.S. & Chesson, P.L. (1989). Diffusion analysis and stationary distribution of the two-species lottery competition model. Theoretical Population Biology, 36, 251–266.

[28] Hatfield, J.S. & Chesson, P.L. (1997). Multispecies lottery competition: a diffusion analysis. In: Structured-Population Models in Marine, Terrestrial, and Freshwater Systems. Springer, pp. 615–622.

[29] Hidalgo, J., Suweis, S. & Maritan, A. (2017). Species coexistence in a neutral dynamics with environmental noise. Journal of theoretical biology, 413, 1–10.

[30] Hoekstra, H.E., Hoekstra, J.M., Berrigan, D., Vignieri, S.N., Hoang, A., Hill, C.E., Beerli, P. & Kingsolver, J.G. (2001). Strength and tempo of directional selection in the wild. Proceedings of the National Academy of Sciences, 98, 9157–9160.

[31] Kalyuzhny, M., Kadmon, R. & Shnerb, N.M. (2015). A neutral theory with environmental stochasticity explains static and dynamic properties of ecological communities. Ecology letters, 18, 572–580.

[32] Kalyuzhny, M., Schreiber, Y., Chocron, R., Flather, C.H., Kadmon, R., Kessler, D.A. & Shnerb, N.M. (2014). Temporal fluctuation scaling in populations and communities. Ecology, 95, 1701–1709.

[33] Kalyuzhny, M., Seri, E., Chocron, R., Flather, C.H., Kadmon, R. & Shnerb, N.M. (2014). Niche versus neutrality: a dynamical analysis. The American Naturalist, 184, 439–446.

[34] Karlin, S. & Liberman, U. (1975). Random temporal variation in selection intensities: one-locus two-allele model. Journal of Mathematical Biology, 2, 1–17.

[35] Karlin, S. & Taylor, H.E. (1981). A second course in stochastic processes. Elsevier.

[36] Kessler, D., Suweis, S., Formentin, M. & Shnerb, N.M. (2015). Neutral dynamics with environmental noise: Age-size statistics and species lifetimes. Physical Review E, 92, 022722.

[37] Kessler, D.A. & Shnerb, N.M. (2007). Extinction rates for fluctuation-induced metastabilities: A real-space wkb approach. Journal of Statistical Physics, 127, 861–886.

[38] Lande, R., Engen, S. & Saether, B.E. (2003). Stochastic population dynamics in ecology and conservation. Oxford University Press.

[39] Leigh, E.G. (2007). Neutral theory: a historical perspective. Journal of Evolutionary Biology, 20, 2075–2091.

[40] Lewontin, R.C. & Cohen, D. (1969). On population growth in a randomly varying environment. Proceedings of the National Academy of sciences, 62, 1056–1060.

[41] Lynch, M. (1987). The consequences of fluctuating selection for isozyme polymorphisms in daphnia. Genetics, 115, 657–669.

[42] Messer, P.W., Ellner, S.P. & Hairston Jr, N.G. (2016). Can population genetics adapt to rapid evolution? Trends in Genetics, 32, 408–418.

[43] Meyer, I. & Shnerb, N.M. (2018). Noise-induced stabilization and fixation in fluctuating environment. Scientific Reports, 8, 9726.

[44] Moran, P.A.P. (1958). Random processes in genetics. In: Mathematical Proceedings of the Cambridge Philosophical Society. Cambridge University Press, vol. 54, pp. 60–71.

[45] Redner, S. (2001). A guide to first-passage processes. Cambridge University Press.

[46] Saccheri, I.J., Rousset, F., Watts, P.C., Brakefield, P.M. & Cook, L.M. (2008). Selection and gene flow on a diminishing cline of melanic peppered moths. Proceedings of the National Academy of Sciences, 105, 16212–16217.

[47] Schreiber, S.J. (2012). Persistence for stochastic difference equations: a mini-review. Journal of Difference Equations and Applications, 18, 1381–1403.

[48] Usinowicz, J., Chang-Yang, C.H., Chen, Y.Y., Clark, J.S., Fletcher, C., Garwood, N.C., Hao, Z., Johnstone, J., Lin, Y., Metz, M.R. et al. (2017). Temporal coexistence mechanisms contribute to the latitudinal gradient in forest diversity. Nature.

[49] Usinowicz, J., Wright, S.J. & Ives, A.R. (2012). Coexistence in tropical forests through asynchronous variation in annual seed production. Ecology, 93, 2073–2084.

[50] Yahalom, Y. & Shnerb, N.M. (2019). Phase diagram for logistic systems under bounded stochasticity. Physical Review Letters, 122, 108102.

[51] Yi, X. & Dean, A.M. (2013). Bounded population sizes, fluctuating selection and the tempo and mode of coexistence. Proceedings of the National Academy of Sciences, p. 201309830.

